# A Passive-Oxygenation Silicone Platform for Biomass Production: Maximizing Labor Productivity and Process Efficiency in Cellular Agriculture Development

**DOI:** 10.64898/2026.06.02.729703

**Authors:** Hiroaki Hatano, Yusaku Takagaki, Misaki Sawada, Ibuki Kokido, Hijiri Okabe, Satoshi Inoue, Kaori Miyaoku, Grace Aprilia Helena, Kota Shiotsuka, Satoshi Tatsumi, Ikko Kawashima

**Author notes:** Equal contribution (Hiroaki Hatano, Yusaku Takagaki).

## Abstract

The commercial production of cell-based food is currently hindered by existing bioreactor technologies, which require substantial capital investment, specialized operating skills, and complex processing setups. To democratize cell-based food production, we developed the “oxy-thru cultivator”—a simple, autoclavable, closed-bag bioreactor fabricated from polydimethylsiloxane (PDMS). By leveraging the high oxygen-permeability of PDMS, this platform enables passive oxygenation across the entire vessel wall, eliminating the need for external aeration or mechanical sparging.

During testing, the cultivator maintained a stable culture environment over 23 days, showing no cytotoxic leachables and retaining both structural integrity and sterility across 10 autoclave cycles. This robustness supported the continuous cultivation of DF-1 cells for 74 days. Using a standardized subculture scheme, we successfully harvested an estimated 2.60 g of cell-based biomass per cultivator over five passages. Notably, the platform achieved a 127% monthly labor productivity compared to conventional bioreactors and was easily operated by researchers without specialized training. Additionally, the system successfully supported the expansion of both mammalian and primary avian cell lines.

With a minimal equipment footprint that reduces CapEx, and a reusable silicone vessel that lowers OpEx, the oxy-thru cultivator offers a highly practical, accessible pathway toward scaling up cellular agriculture.

**Graphical Abstract:** 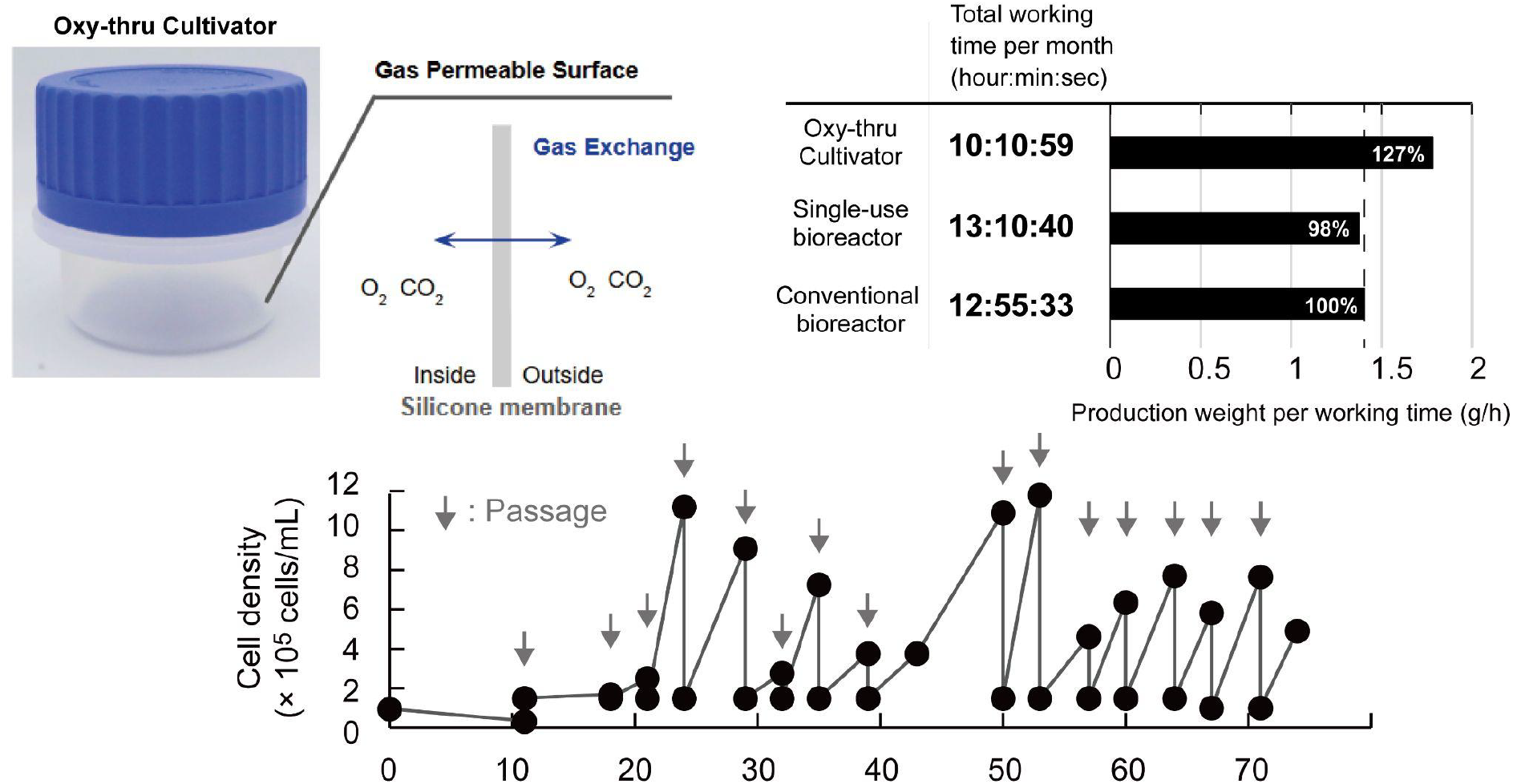

## 1. Introduction

Achieving efficient and scalable cell culture is fundamental to productivity and cost-effectiveness in tissue engineering, regenerative medicine, and the rapidly emerging field of cellular agriculture,(1–3) yet the industrialization of cellular agriculture in particular remains constrained by significant barriers that limit participation to well-resourced organizations.(4,5) Advancing the field requires not only scientific progress but also the democratization of biotechnology—broadening participation from academic institutions and startups to individual innovators.(6–9) To achieve this, it is essential for each stakeholder to establish a minimum capacity for cell-based biomass production without prohibitive capital investment.(10,11)

From a technical perspective, the primary obstacle to establishing this production baseline is the inherent difficulty of managing gas exchange as culture volumes expand. In existing bioreactor technologies, achieving sufficient dissolved oxygen (DO) levels typically requires substantial capital investment, specialized operating skills, and high process complexity. (12–14) Furthermore, as culture volume increases, the efficient delivery of dissolved oxygen (DO) becomes a critical challenge,(15,16) typically addressed through mechanical agitation or perfusion systems,(17–20) approaches that further increase system complexity and introduce the risk of shear-induced cell damage.(21,22) Consequently, there is a pressing need for simple, cost-effective, and cell-friendly culture systems capable of supporting biomass production across diverse cell types without requiring specialized infrastructure or expertise.(23,24)

To overcome these limitations, we focused on silicone, especially polydimethylsiloxane (PDMS), which is widely used as a biomaterial.(25–27) In addition to excellent biocompatibility and processability,(28,29) PDMS possesses exceptionally high permeability to gases, particularly oxygen, compared to many other polymeric materials.(30–32) Leveraging the inherent high oxygen permeability of PDMS, we hypothesized that sufficient oxygen could be passively transferred from the ambient air through the vessel wall into the culture medium, thereby eliminating the need for external aeration ports or mechanical agitation. Based on this concept, we designed a closed culture bag with a very simple structure composed of silicone—the “oxy-thru cultivator”—comprising only three components, and validated its compatibility with FDA 21 CFR 177.2600 for food-contact applications.

The objective of this study was to evaluate whether the oxy-thru cultivator could support stable cell culture without additional oxygen supply, and to assess its practical advantages in terms of operational efficiency, infrastructure requirements, and versatility across diverse cell types relevant to cellular agriculture applications.

## 2. Materials and Methods

### 2.1. Fabrication and Characterization of oxy-thru cultivator

The oxy-thru cultivator is a modular, passive-oxygenation bioreactor fabricated via injection molding using medical-grade polydimethylsiloxane (PDMS) silicone rubber (KE1950; Shin-Etsu Chemical Co., Ltd., Tokyo, Japan). To ensure structural stability and a low center of gravity while maintaining high gas permeability, the vessel was designed with a specific wall thickness gradient. The side walls range in thickness from 1.0–1.6 mm at the opening (thin-walled section) to 2.4–7.8 mm near the base (thick-walled section), with an inclination angle of 5° from the vertical axis. The vessel body is integrated with a port assembly comprising a polypropylene (PP) lid (GL-50; German Industrial Standard DIN 168) and a cap holder, both capable of withstanding high-pressure steam sterilization. The assembly ensures a liquid-tight seal through a screw-thread mechanism that compresses a flange-like protrusion on the silicone vessel. Prior to use, all components were sterilized by autoclaving at 121°C for 20 min.

### 2.2. Oxygen Measurement and Environmental Stability

To evaluate the oxygen supply capacity of the oxy-thru cultivator, gas permeation properties and culture environment parameters were characterized. The oxygen permeability of the PDMS membrane was measured by an external testing laboratory (Aichi Center for Industry and Science Technology; Aichi Japan) using a gas permeability analyzer (GP-1000AZ; J-SCIENCE LAB CO.,Ltd., Kyoto, Japan). For comparative analysis, two other vessels made of polyethylene (PE) were used: a commercial culture bag A (87-352, Nipro Corporation Osaka, Japan), and a commercial culture bag B (FKCB215, Fukoku Co., Ltd., Saitama, Japan).

For the in-vessel measurement of gas partial pressures and dissolved oxygen (DO), the cultivators were placed in a standard incubator at 37°C with 5% CO_2_ and 60% humidity. Cultures were subjected to constant 8-figure orbital shaking at 60 rpm to simulate operational conditions. After 23 days of continuous operation, key parameters including pH, oxygen partial pressure O_2_, and carbon dioxide partial pressure CO_2_ were quantified using a blood gas analyzer (GASTAT-720; Techno Medica Co., Ltd., Kanagawa, Japan). Dissolved O_2_ concentration (mg/L) was subsequently derived from the partial pressure values.

### 2.3. Structural Robustness and Safety Validation

The dimensional stability of the silicone vessel was assessed across 1 to 10 autoclave cycles. Sterility was confirmed via the absence of colony-forming units (0 CFU) on Soybean-Casein Digest (SCD) agar and incubated at 35°C. Colony-forming units (CFU) were quantified using an automatic colony counter (Scan500; Interscience, Saint Nom la Bretêche, France). For safety validation (FDA 21 CFR 177.2600), elution tests were conducted under various solvent conditions to detect hazardous leachables, including heavy metals and phenolic compounds. These safety evaluations were commissioned and performed at the Japan Food Research Laboratories (Mie, Japan) to ensure the vessel’s suitability for repeated food-contact applications.

### 2.4. Cells and Cell Culture

The chicken embryonic fibroblasts UMNSAH/DF-1 cell line (DF-1 cell) was purchased from ATCC (CRL-3586, Virginia, USA). Immortalized porcine skeletal muscle satellite cell line was purchased from Creative Bioarray (CBADE0935, NY, USA). The mouse myoblast cell line (C2C12 cell) was purchased from RIKEN (RCB0987, Ibaraki, Japan). Primary duck liver-derived cells were isolated and cultured as previously described.(33)

All mammalian cells were maintained in Dulbecco’s Modified Eagle Medium (DMEM; Gibco, Waltham, MA, USA) with 10% Fetal Bovine Serum (FBS; Gibco, Waltham, MA, USA and 1% Penicillin-Streptomycin-Amphotericin B (PSA; Fuji-film-Wako, 161-23181). All avian cells were maintained in food-grade D-MEM replacement I-MEM1.0 (IntegriCulture Inc., Tokyo, Japan) (34) supplemented with 10% a yolk-derived lipoprotein (YLP) serum alternative supplement (35) and 1% PSA. Prior to seeding, the culture surfaces for primary duck liver-derived cells were coated with iCoater (Integriculture Inc., Tokyo, Japan) and cultured as previously described.(36) Cultures were maintained in a standard incubator at 37°C with 5% CO_2_. For each passage, cells were detached using a control reagent (TrypLE™ Express; 12605010; Thermo Fisher Scientific, MA, USA) or iDisper, a food-grade cell dissociation agent (IntegriCulture, Tokyo, Japan) as previously described,(33) for 10 min incubation at 37 °C and 60% humidity. The cell suspension was centrifuged at 300 × g for 4 min at room temperature, and the pellet was pooled and seeded into a single T175 flask to minimize individual variability.

### 2.5. Suspension and Scaffold Cell Culture

Suspension cultures were performed in the oxy-thru cultivator (200 mL working volume) and the cultistar (30 mL and 100 mL scales; ABLE Corporation, Tokyo, Japan). The cultistar was operated on a multi-point stirrer (BWS-S03N0S-6C; ABLE Corporation, Tokyo, Japan) at 40 rpm. The oxy-thru cultivator was equipped with a magnetic stir bar (diameter: 30 mm; KOKUGO Co., Ltd., Tokyo, Japan) and operated on a magnetic stirrer (40101; TOHO KK., Tokyo, Japan) at 40 rpm.

For experiments using cell attachment scaffolds conducted in the oxy-thru cultivator, a gelatin-based scaffold was used in our previous patent.(37) These cultures were agitated at 60 rpm using an orbital shaker (In-Vitro Shaker 2 Shake-XR2; TIETECH Co., Ltd., Nagoya, Japan).

### 2.6. Cell Count and Metabolite Measurement

The cell number and viability were measured using an automated cell counter (NucleoCounter® NC-202™; ChemoMetec, Denmark) with Via2-Cassette (941−0024; ChemoMetec, Denmark) containing acridine orange and 4′,6-diamidino-2-phenylindole (DAPI) fluorescent dyes, according to the manufacturer’s instructions. Due to the aggregation of DF-1 cells in suspension, samples were treated with TrypLE™ Express for 10 min followed by vigorous pipetting prior to counting; thus, “total cells” was used as the indicator. For scaffold-based cultures, 5 mL of iDisper + EDTA solution (311-90075; Fujifilm Wako, Tokyo, Japan) was added to the tube and incubated in a water bath at 37°C for 30 min to dissolve the scaffold and detach cells. The resulting solution was pipetted with a 1,000 µL pipette, and 1 mL of the suspension was transferred to a 1.5 mL tube for counting.

Glucose and lactate concentrations were measured using a multiple biosensor (BF-9M; Oji Scientific Instruments, Tokyo, Japan).

### 2.7. Operating Time Measurement

Labor time measurements were initiated once all consumables and equipment were prepared. The recorded time represents the actual working time of the operator and excludes autoclave processing time. Fixed time values were assigned for tasks ancillary to cell counting: centrifugation (00:05:25), supernatant disposal (00:02:50), and cell counter measurement (00:05:00). All other tasks were measured in real-time. The “Researcher” had more than three years of experience in standard 2D cell culture, while the “Expert” possessed over five years of experience in bioreactor operations. Since the conventional bioreactor inherently requires expert-level operation due to its structural complexity—including the precise assembly of multiple tube joints, and functional ports—the assignment of an expert operator to that system reflects its actual deployment condition rather than an experimental bias. Accordingly, this comparison represents an operationally realistic scenario for each system, and the difference in operator skill level is itself a measurable characteristic of the respective platforms.

The conventional bioreactor system used for comparison consisted of a 1,000–5,000 mL glass vessel bioreactor (BCP-S05NP 4S, ABLE Corporation, Tokyo, Japan) and its control system. The assembly of this system requires the precise installation of numerous discrete components, including multiple O-rings, various tube joints, and functional ports (Supplementary Figure 1). Due to this high level of structural and operational complexity, which involves significant risks of contamination or leakage if improperly handled, the participation of an “Expert” operator was essential to ensure reproducible and sterile setup. Other equipment included a standard incubator (MCO-80IC-PJ; PHC Corporation, Tokyo, Japan), a laminar flow hood (MCV-161BNS-PJ; PHC Corporation, Tokyo, Japan), an autoclave(HV-110 II LB; HIRAYAMA Manufacturing Corporation, Satitama, Japan), an oxygen generator (02-PSA MOX-0.8 II; KOFLOC, Kyoto, Japan), a peristaltic pump (77201-62, MASTERflex L/S; Yamato Scientific co., ltd., Tokyo, Japan), a tube sealer (16391-000, BIOSEALER TC, SARTORIUS; Göttingen, Germany), and a tube welder (BWTC10061, BIOSEALER TC, SARTORIUS; Göttingen, Germany).

### 2.8. Productivity Analysis

#### 2.8.1. Biomass Production Calculation

Cumulative biomass production was calculated based on the growth kinetics of DF-1 cells shown in Figure 3E. Subculturing and cell recovery were triggered whenever the cell density exceeded a passage threshold of 4.5×10^5^ cells/mL. At each passage, cells were re-seeded at a density of 1.0-1.5×10^5^ cells/mL, and the remaining surplus cells were defined as the recovered biomass. The cumulative biomass was calculated for both a single-unit (1x) and a five-unit (5x) oxy-thru cultivator system. According to the growth profile, the recovery cycles were set on Day 60 (1st), Day 64 (2nd), Day 67 (3rd), Day 71 (4th), and Day 74 (5th).

#### 2.8.2. Labor Productivity Calculation

Labor productivity (g/h) was calculated to evaluate the industrial efficiency of each system. The monthly cell-based biomass yield was estimated by multiplying the mean number of cells recovered per passage (based on the five passages shown in Figure 3G) by the total number of passages per month (7 passages, Figure 4C). The total monthly labor time was determined by summing the real-time measurements for these 7 cycles. Finally, labor productivity was derived by dividing the total monthly biomass (calculated using the dry cell weight of 5 ng/cell) by the total monthly labor time.

### 2.9. Statistical Analysis

All experiments were conducted with at least three biological replicates (*n*=3). Data are presented as the mean ± standard deviation (SD). Statistical significance between two experimental groups was determined using a two-tailed Student’s t-test. A *p*-value of less than 0.05 was considered statistically significant.

## 3. Results

### 3.1. Design of the “oxy-thru cultivator”

We developed a passive-oxygenation bioreactor, termed the “oxy-thru cultivator,” designed to support cell growth via gas permeation through a silicone membrane. This system eliminates the requirement for external gas supply infrastructure and specialized attachments (Figure 1). The oxy-thru cultivator comprises three primary components: a silicone vessel serving as the culture area, a lid, and a bearing joint that secures the assembly. The vessel is fabricated from a biocompatible silicone sheet, and the port assembly lid ensures an airtight seal while allowing convenient access for media exchange and cell seeding. Owing to its structural and operational simplicity, the oxy-thru cultivator enables researchers to conduct 3D cell culture through simple assembly and routine laboratory procedures.

**Figure 1:**
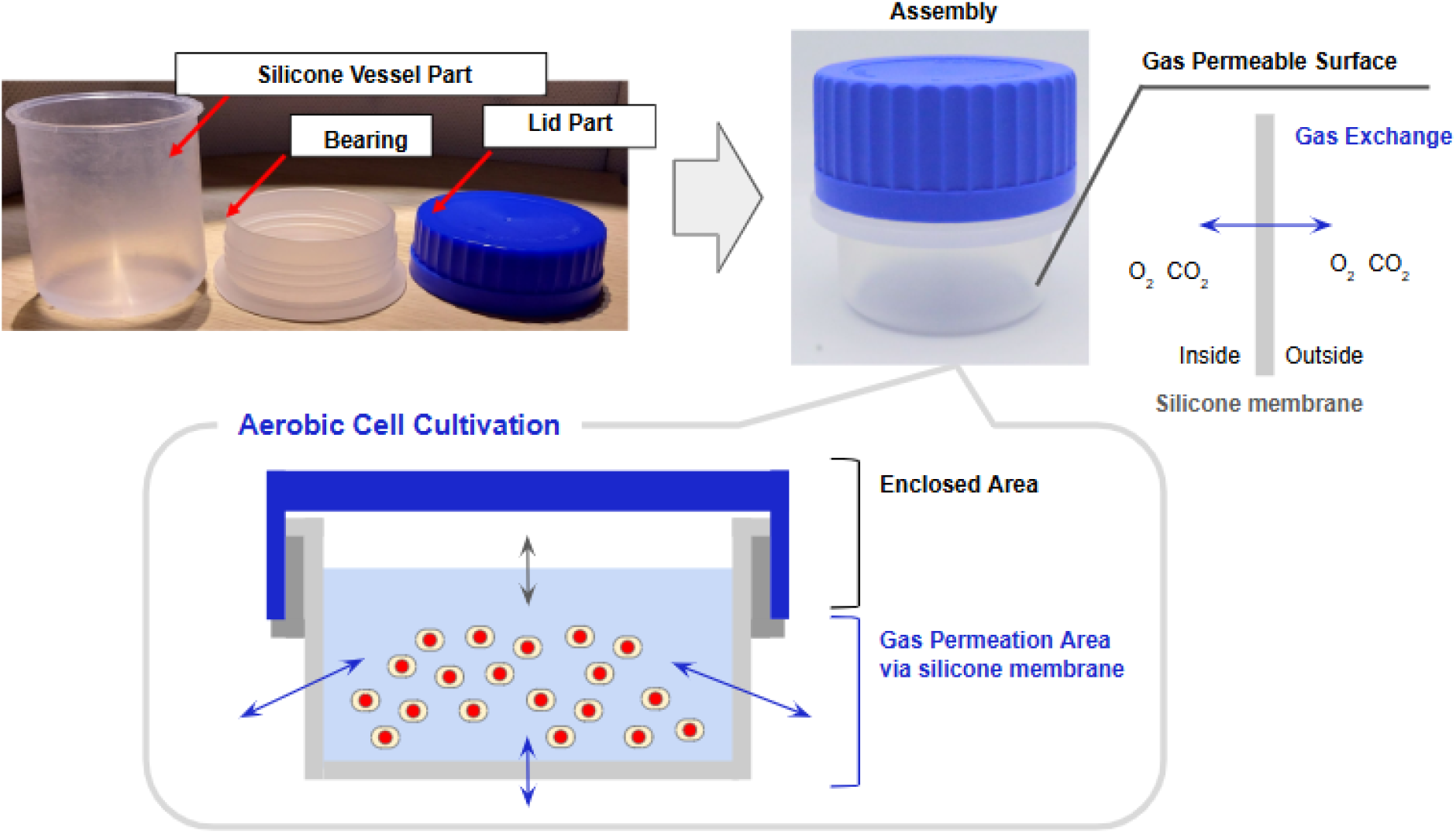
Components and operating principle of the oxy-thru cultivator. The figure displays the individual components, the assembled vessel, and the conceptual mechanism of the cultivation system. The cultivator comprises three primary parts: a silicone vessel (culture area), a lid, and a bearing joint. The schematic diagram illustrates the principle of passive oxygenation, where ambient gases undergo permeation through the silicone membrane. This design facilitates efficient gas exchange across the entire surface of the gas-permeable vessel, enabling cell culture without the need for external gas supply systems.

### 3.2. Silicone Material Properties: Gas Permeability and Safety Validation

The oxygen permeability and biocompatibility of the silicone material were validated to ensure its suitability for food-related applications, specifically following FDA 21 CFR 177.2600 regulations for repeated food-contact use.

First, we characterized the gas permeation properties of the silicone membrane. Despite being significantly thicker (0.5–1.0 mm vs. 0.1 mm), the silicone sheets exhibited oxygen permeability one to two orders of magnitude higher than that of conventional polyethylene-based culture vessels, demonstrating substantially superior potential for passive gas exchange (Figure 2A). To evaluate the impact on the culture environment, key parameters within the oxy-thru cultivator were compared against a commercial culture bag and a glass bottle after 23 days of operation. The oxy-thru cultivator maintained higher dissolved oxygen (DO) levels (5.5 mg/L) and a more stable pH (7.43), values notably more favorable than those observed in the commercial bag and glass bottle controls (Table 1).

**Table 1.** Comparison of culture environment parameters across different vessel types on Day 23. Initial medium values are provided as a baseline to demonstrate the stability of pH and gas partial pressures (O_2_ and CO_2_) in the oxy-thru cultivator compared to commercial bags and glass bottles.

| Parameter | Unit | Initial Medium | oxy-thru cultivator<br>(Day 23) | Commercial Bag<br>(Day 23) | Glass bottle<br>(Day 23) |
| --- | --- | --- | --- | --- | --- |
| pH | - | 7.71 | 7.43 | 7.28 | 7.69 |
| O<sub>2</sub> Partial Pressure | mmHg | 153.1 | 122.1 | 116.5 | 99.9 |
| CO<sub>2</sub> Partial Pressure | mmHg | 31.8 | 40.4 | 53.9 | 16.6 |
| Dissolved O<sub>2</sub> | mg/L | 6.9 | 5.5 | 5.3 | 4.5 |
| Dissolved CO<sub>2</sub> | mg/L | 46.9 | 59.5 | 79.4 | 24.5 |

**Figure 2:**
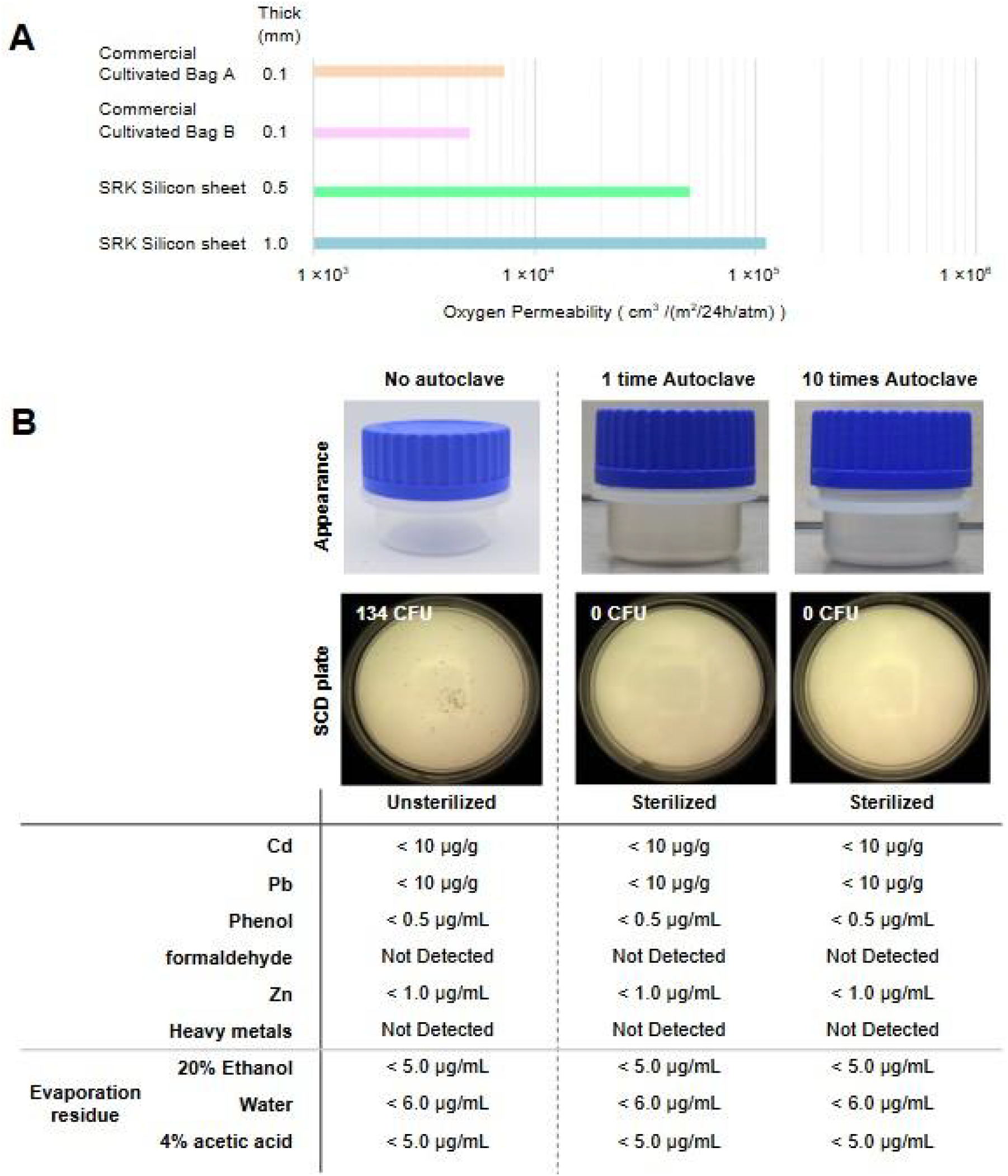
Structural stability and chemical safety of the oxy-thru cultivator. (A) Comparison of oxygen permeability between silicone membranes (0.5 mm and 1.0 mm) and commercial polyethylene-based culture bags (0.1 mm). (B) Validation of autoclave stability and material safety. Top: Visual appearance and sterility (SCD plate) after repeated autoclave cycles. Bottom: Elution test results demonstrating compliance with safety standards for food-contact materials.

**Figure 3.**
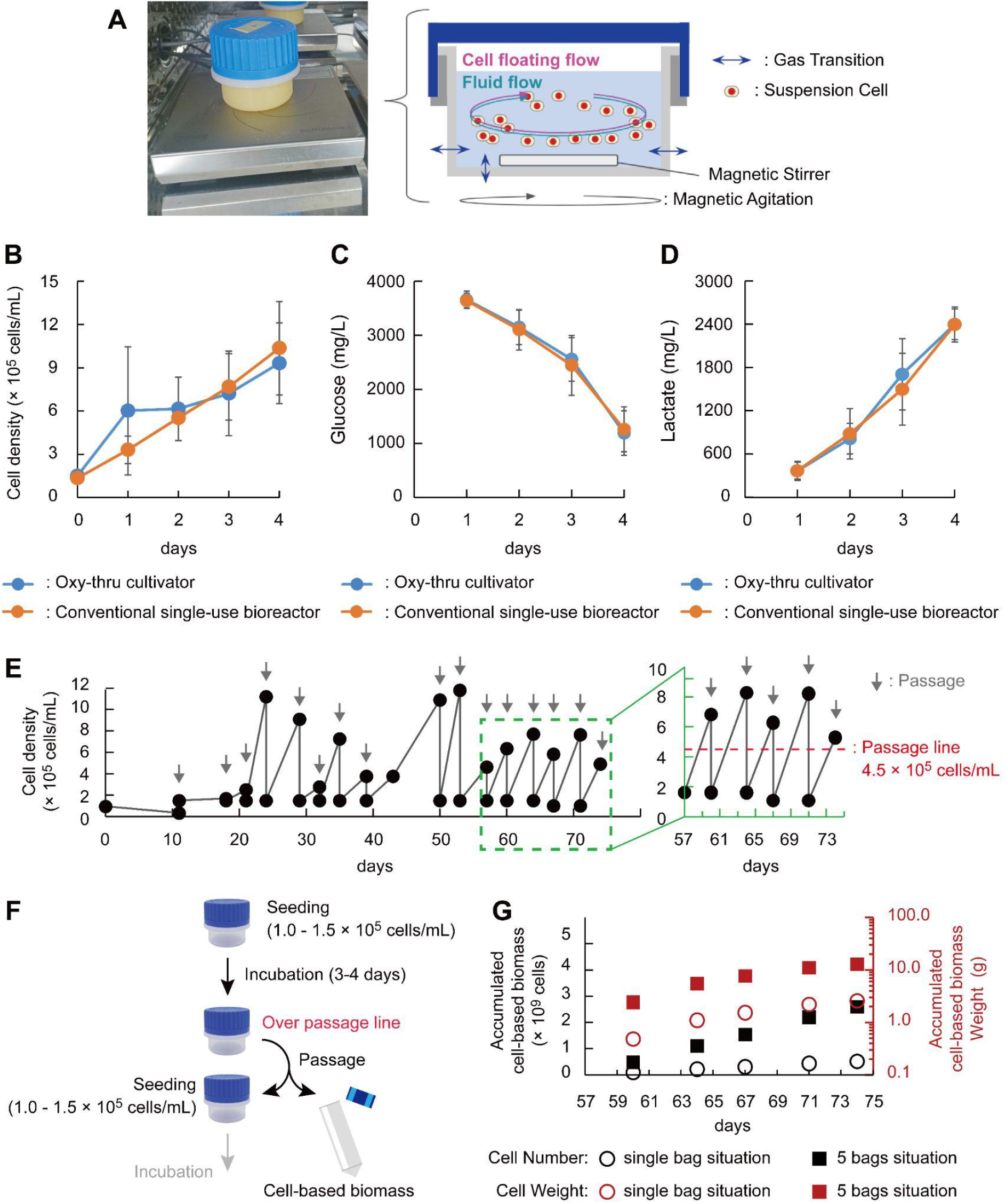
Evaluation of long-term suspension culture and biomass production in the oxy-thru cultivator. (A) Experimental setup for agitated culture using a magnetic stirrer. (B–D) Temporal changes in cell density, glucose concentration, and lactate concentration. Data are presented as mean ± SD (*n*=3).The single-use (SU) bioreactor used in this comparison was cultistar. No significant differences were observed between the oxy-thru cultivator and cultistar across all parameters throughout the culture period. (E) Representative growth curve of DF-1 cells during long-term continuous passage (74 days). Gray downward arrows indicate the timing of cell passages. The red dotted line represents the Passage Line (4.5 ×10^5^ cells/mL); the period during which this Passage Line was determined and fixed is indicated by the green box.(F) Subculture process flow. Passage is performed once the cell density exceeds the Passage Line; cells not utilized for seeding are collected and stored separately as cell-based biomass. (G) Calculated accumulated cell-based biomass. The black Y-axis (left) indicates the accumulated cell number, and the red Y-axis (right) indicates the accumulated weight. Symbols represent a single unit (〇) and five units (▢) of the oxy-thru cultivator.

**Figure 4.**
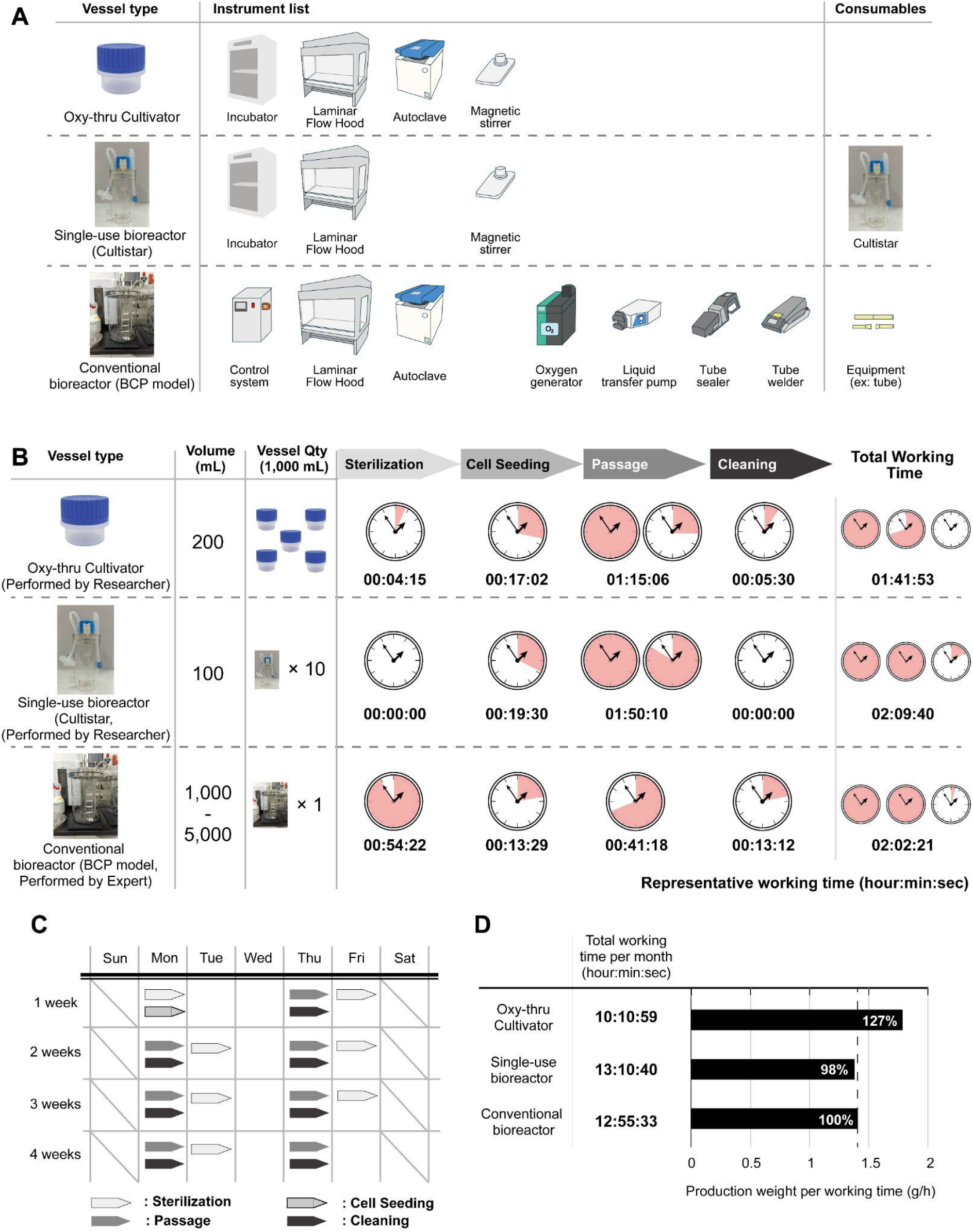
Comparison of resource requirements, operational efficiency, and labor productivity. (A) List of required instruments and consumables for the Oxy-thru cultivator, single-use bioreactor, and conventional bioreactor. (B) Labor time for sterilization, cell seeding, passage, and cleaning at a 1,000 mL working volume. The Oxy-thru and single-use systems were operated by a general researcher, while the conventional bioreactor was operated by an expert (>5 years experience). (C) Calculated a 1 month (4 weeks) production calendar based on seven passages. (D) Total monthly labor time and labor productivity (produced weight per working time, g/h). The productivity of the conventional bioreactor is defined as 100% (dotted line).

Next, the structural robustness and sterility of the vessel were assessed. The oxy-thru cultivator maintained dimensional stability and integrity after 1 to 10 autoclave cycles, with sterility confirmed by the absence of colony-forming units (0 CFU) on SCD plates (Figure 2B). Furthermore, elution tests revealed that no hazardous or cytotoxic substances, including heavy metals and phenolic compounds, leached from the vessel under various solvent conditions (Figure 2B). These results confirm that the vessel provides a robust, sterile, and chemically safe environment for long-term cell culture.

### 3.3. LONG-term Stability and Standardization of Cell Proliferation in Agitated Culture

To evaluate the performance of the oxy-thru cultivator in cellular agriculture applications, DF-1 cells (chicken embryonic fibroblasts) were employed as a model. DF-1 cells are an industrially important cell line, notably utilized by GOOD Meat Inc. for the first FDA-approved cultured meat product in 2022. (38) For suspension culture, the oxy-thru cultivator was equipped with a magnetic stir bar and placed on a magnetic stirrer inside a standard incubator (Figure 3A).

The growth kinetics and metabolic profiles in the oxy-thru cultivator were compared against a conventional single-use bioreactor (cultistar). The oxy-thru cultivator exhibited cell growth trends comparable to the conventional system (Figure 3B). Furthermore, glucose consumption (Figure 3C) and lactate production (Figure 3D) profiles showed no significant difference between the two systems. This demonstrates that the oxy-thru cultivator provides an environment for suspension culture equivalent to existing commercial bioreactors.

We further validated the feasibility of long-term continuous cultivation. Following recovery from cryopreservation, DF-1 cells were successfully maintained in the oxy-thru cultivator for 74 consecutive days (Figure 3E). From Day 57, the subculture interval was standardized to a 3-4 day cycle with a fixed passage threshold. This result demonstrates that the oxy-thru cultivator supports a highly consistent and predictable expansion protocol for avian suspension cells over an extended period.

Toward the practical application of cell-based biomass production, a recovery scheme was established by standardizing the passage threshold and seeding density (Figure 3F). Based on this scheme, the cumulative biomass production was calculated for the period between Day 57 and Day 74 (Figure 3G). For the biomass calculation, the weight of a single DF-1 cell was defined as 5.0 ng/cell, based on experimentally measured cell pellet weights. A dry cell weight of 5.0 ng/cell was derived from experimentally measured wet weights of DF-1 cell pellets (63.0 ± 9.7 ng/cell; n=4) after correction for moisture content (92.13%), as previously reported for duck liver-derived cell-based food.(39) The large standard deviation reflects biological variability in pellet preparation and represents a recognized source of uncertainty in the biomass estimate. The measurements indicated that each oxy-thru cultivator yields approximately 0.52 ± 0.12 g of cell-based biomass per passage (*n*=5). By the final day (after 5 passages), a total of 2.60 g would be recovered. Furthermore, if five oxy-thru cultivators were used, the recovered cell weight at Day 3 (1 passage) would be equivalent to that of a single unit, reaching a total of 12.99 g by Day 17.

### 3.4. Comparative Analysis of Resource Requirements and Labor Productivity

The oxy-thru cultivator is autoclavable (Figure 2B) and facilitates cell culture within a standard incubator, comparable to conventional single-use systems (Figures 3B–D). In contrast, conventional non-single-use bioreactors (e.g., jar fermenters) require extensive peripheral infrastructure, including external control systems, oxygen generators, and specialized equipment for sterile tube welding and sealing. Figure 4A summarizes the required instruments and consumables for each system. The minimized instrument count for the oxy-thru cultivator directly reduces Capital Expenditure (CapEx), while the reusable silicone vessel notably lowers Operational Expenditure (OpEx) compared to single-use bioreactors, which require continuous replacement of specialized consumables.

Operational efficiency was evaluated by measuring the labor time for four essential culture processes at a 1,000 mL working volume (Figure 4B). For a rigorous comparison, the oxy-thru cultivator was operated by a general researcher, while the conventional bioreactor was handled by an expert with over five years of experience due to its system complexity. In the oxy-thru cultivator and single-use systems, “Passage” was the most labor-intensive step. Conversely, for the conventional bioreactor, the assembly and sterilization of numerous components required the longest working time. Overall, the oxy-thru cultivator required the least total labor time among the three systems.

Based on the subculture scheme (Figure 3F) and growth kinetics (Figure 3E), a 30-day production calendar was calculated (Figure 4C). Assuming seven recovery passages per month—accounting for standard weekends and shift-based task rotations—the total monthly labor time and biomass yield (derived from Figure 3G) were integrated to calculate labor productivity (Figure 4D). The oxy-thru cultivator achieved a monthly labor productivity of 127% compared to the conventional bioreactor, demonstrating that the system enables high-efficiency production even when operated by researchers without specialized expertise.

While mass production for cell-based food research is often limited by prohibitive CapEx and labor intensity, the oxy-thru cultivator enables 1,000 mL-scale biomass production using standard laboratory equipment. This system provides a highly efficient and practical platform for cellular agriculture without the need for specialized infrastructure or unsustainable labor schedules.

### 3.5. Versatile Application of the oxy-thru cultivator

To evaluate the versatility of the oxy-thru cultivator, we conducted culture tests using various cell types. Given that most cells in cellular agriculture are adherent, we implemented an adherent culture system by placing scaffolds within the oxy-thru cultivator. The vessel was then positioned on a shaker to ensure medium uniformity through gentle agitation (Figure 5A).

**Figure 5.**
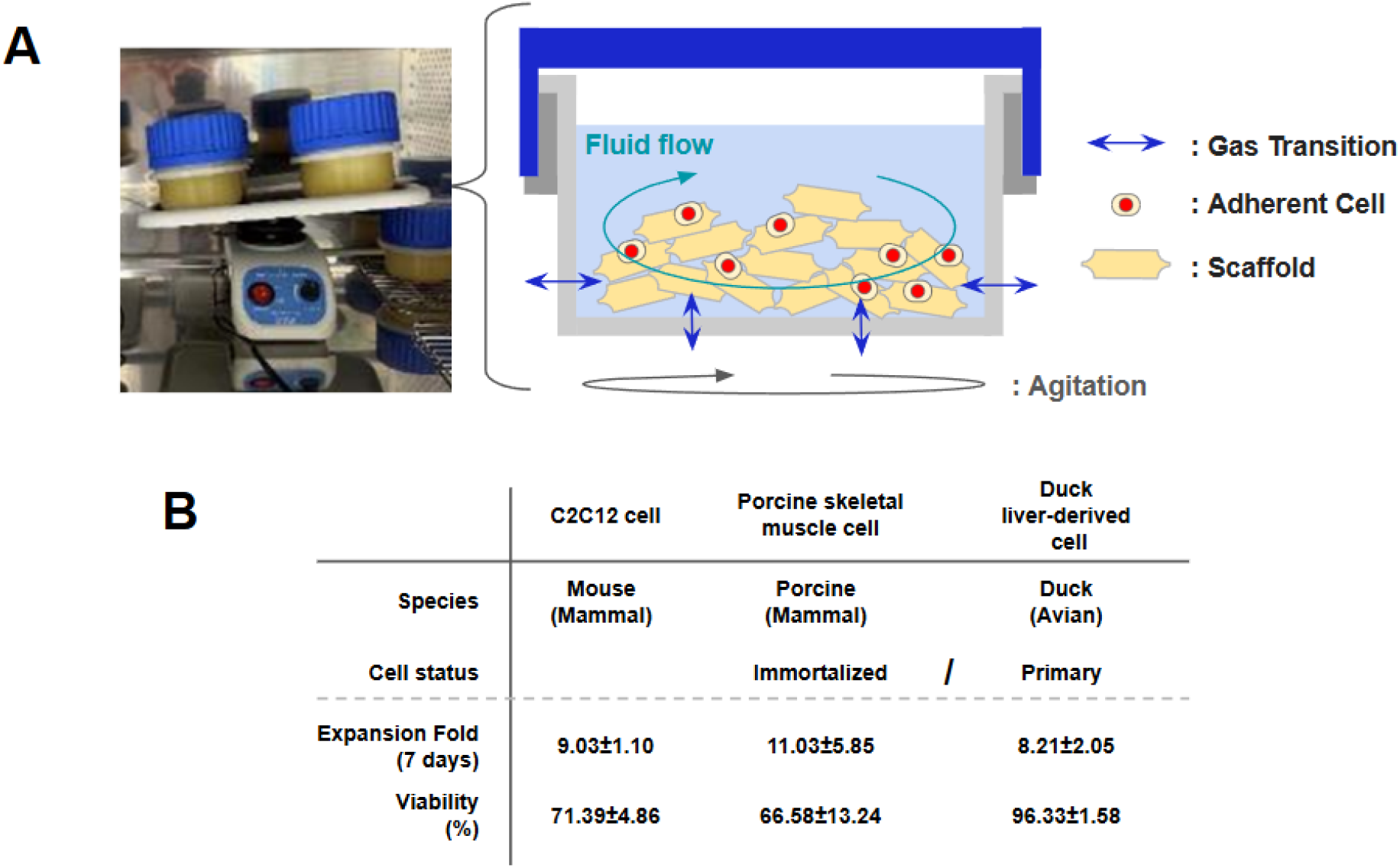
Versatility of the oxy-thru cultivator across different animal species and cell types. (A) Schematic and photographic representation of the adherent culture setup using scaffolds and orbital agitation. (B) Summary of expansion fold (at 7 days) and cell viability for mouse (C2C12), porcine (skeletal muscle), and duck (liver-derived primary) cells. Data are presented as mean ± SD (*n*=4), representing the combined results from both static and agitated culture conditions.

Under these conditions, we confirmed the successful expansion of several cell lines over a 7-day period, including mammalian cells (mouse C2C12 and porcine skeletal muscle satellite cells) and avian cells (primary duck liver-derived cells). All tested cell types—mammalian and primary avian—maintained a viability exceeding 60% (Figure 5B); notably, duck liver-derived primary cells achieved 96.3%, suggesting particularly strong compatibility with the platform. Since all cell types were cultured under identical conditions in this study, further optimization of culture parameters for each cell type is expected to improve viability and expansion efficiency. Nevertheless, these results demonstrate that the oxy-thru cultivator provides a versatile platform capable of supporting cell growth across a wide range of animal species and cell types, overcoming the barriers between mammalian and avian cell culture.

## 4. Discussion

The oxy-thru cultivator leverages the high gas permeability of PDMS to enable passive oxygenation across the entire vessel wall, rather than restricting gas exchange to the medium-surface interface. In addition, the oxygen permeability of the silicone membrane was orders of magnitude higher than that of conventional polyethylene-based culture vessels. This property maintained a stable culture environment at Day 23, with an O_2_ partial pressure of 122.1 mmHg—significantly higher than that of the glass bottle (99.9 mmHg)—and a stable pH of 7.43 (Table 1). Importantly, this passive aeration eliminates mechanical sparging—a primary source of shear-induced cell damage.(21,40,41) This stable culture environment supported continuous DF-1 cell cultivation for 74 days, with a standardized subculture scheme yielding an estimated 2.60 g of cell-based biomass per cultivator over 5 passages, demonstrating the system’s suitability for long-term and quantifiable biomass production (Figures 3E, 3G). The practical consequence of this design simplicity was substantial: sterilization required only 4 min 15 sec (00:04:15) by a general researcher, versus 54 min 22 sec (00:54:22) by an expert operator for a conventional bioreactor, yielding a monthly labor productivity of 127% relative to conventional systems (Figure 4D). Notably, this comparison is inherently conservative: the conventional bioreactor was operated by an expert with over five years of experience, whereas the oxy-thru cultivator was operated by a researcher without specialized bioreactor training. The significance of this asymmetry becomes clear when one considers that conventional bioreactor operation is widely recognized as a distinct technical discipline requiring dedicated education. Multiple studies have independently developed structured curricula and low-cost platforms specifically to lower the expertise barrier associated with bioreactor setup and operation—a need that itself attests to the non-trivial expertise barrier that conventional bioreactor operation imposes.(42–45) Furthermore, the prohibitive cost of conventional systems, often ranging from $10,000 to $40,000 per unit, compounds this barrier, restricting access to well-resourced institutions.(45) When the hidden costs of expert recruitment, long-term training investment, and personnel retention are taken into account, the true operational advantage of the oxy-thru cultivator is likely to be substantially greater than the measured 127% figure suggests. Consequently, the system can be operated at production scale by personnel with basic laboratory training, without the need for dedicated specialists or extensive operator development programs.

In terms of infrastructure, the oxy-thru cultivator requires only four standard instruments—incubator, laminar flow hood, autoclave, and magnetic stirrer—with no dedicated consumables, in contrast to conventional bioreactors that additionally require oxygen generators, liquid transfer pumps, tube sealers, and tube welders (Figure 4A). This minimal equipment footprint directly reduces CapEx, while the reusable silicone vessel eliminates the recurring consumable costs inherent to single-use systems, lowering OpEx. Furthermore, because the PDMS material is pre-validated under FDA 21 CFR 177.2600 and elution testing confirmed the absence of cytotoxic leachables across repeated autoclave cycles (Figure 2B), the regulatory hurdles associated with material safety are largely resolved at the platform level, further reducing the effective cost of entry into cellular agriculture production.

The platform’s applicability across mammalian and avian cell types, including primary cells, further demonstrates that these advantages are not limited to a specific cell line, broadening its potential user base across the diverse landscape of cellular agriculture.

Together, these properties address the critical barriers to industrialization in cellular agriculture—cost, complexity, and operational and regulatory burden—by leveraging the high gas permeability of PDMS.

Although the current platform was validated at a 200 mL working volume, the relationship between vessel surface area, membrane thickness, and oxygen demand at larger scales warrants further investigation, representing a natural next step toward commercial-scale implementation.

## 5. Conclusion

The oxy-thru cultivator demonstrates that the high oxygen permeability of PDMS alone can sustain a stable culture environment sufficient for long-term cell culture, as evidenced by continuous DF-1 cell cultivation for 74 days. Furthermore, a standardized subculture scheme enabled quantifiable biomass recovery of 2.60 g per cultivator over 5 passages, providing a concrete production benchmark for cellular agriculture research. Beyond this, the system’s applicability across mammalian and avian cell types, including primary cells, confirms its versatility as a platform for the diverse cell lines relevant to cellular agriculture. By requiring only standard laboratory instruments, eliminating specialized consumables, and leveraging a pre-validated food-contact material, the oxy-thru cultivator simultaneously reduces CapEx, OpEx, and regulatory burden. Together, these properties lower the barriers to entry that have historically confined cell-based biomass production to well-resourced organizations, offering a practical and accessible platform for the broader industrialization of cellular agriculture.

## Data Availability Statement

The datasets generated and analyzed during the current study are not publicly available due to proprietary and commercial confidentiality but may be available from the corresponding author on reasonable request.

Key results are summarized in the main text, with additional supporting data available in the supplementary files.

## Author Contributions

H.H. designed the study and experiments. Y.T. conceived the development concept. Experiments were conducted by Y.T., M.S., I.Ko., H.O., S.I., G.A.H., and K.S. Data interpretation and analysis were performed by H.H., Y.T., and K.M. The original draft was written by H.H. and reviewed and edited by H.H. and I.Ka. H.H., S.T., and I.Ka. provided overall supervision and authorized the study. All authors have read and approved the final manuscript.

## Acknowledgments

We are deeply grateful to Mr. Naoki Funamoto of Sumitomo Riko Co., Ltd., who played a pivotal role in establishing this research collaboration. By bridging our respective organizations, he made this entire project possible, and we sincerely thank him for his foundational contributions.

