## Supplementary Materials for "A Passive-Oxygenation Silicone Platform for Biomass Production: Maximizing Labor Productivity and Process Efficiency in Cellular Agriculture Development"

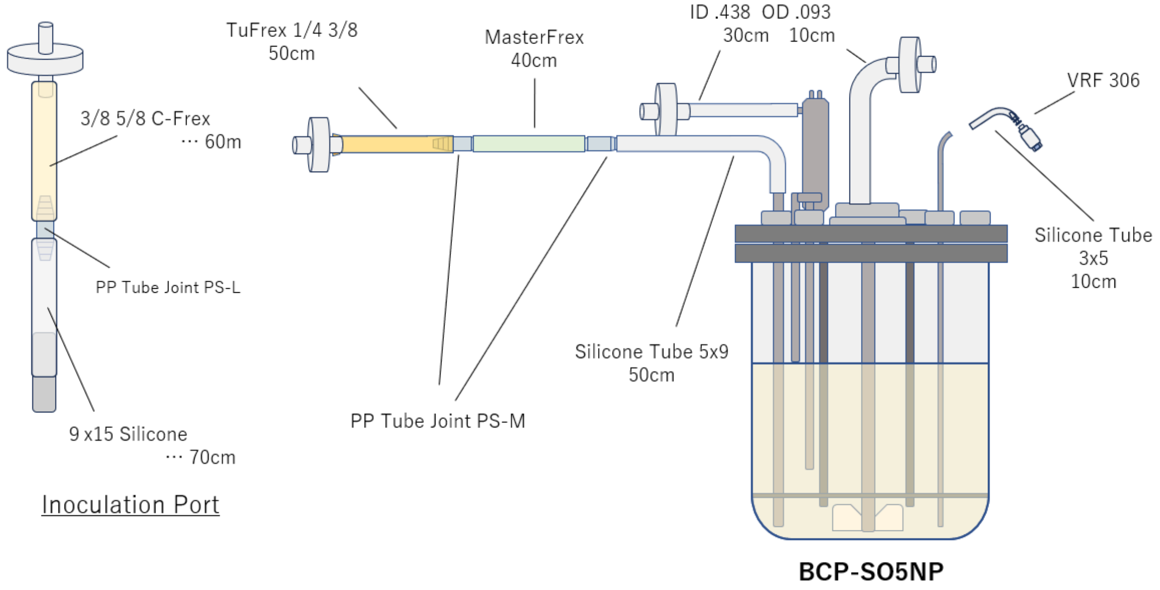


**Supplementary Figure 1: Schematic representation of the BCP-SO5NP 4S bioreactor assembly.** The illustration highlights the significant operational complexity involved in the preparation of a conventional stirred-tank bioreactor. The system requires the integration of over a dozen specialized components, including an inoculation port with various tube connectors (e.g., PP Tube Joint PS-L/PS-M), C-Flex and silicone tubing of varying diameters, and a headplate assembly.
